# Transient dynamics and counterintuitive competitive performance in periodic environments

**DOI:** 10.1101/2025.10.26.684656

**Authors:** Carling Bieg, Bailey C McMeans, Alexa Scott, Kevin S McCann

**Affiliations:** Department of Integrative Biology, University of Guelph, Guelph, Ontario, Canada; Department of Biology, University of Toronto Mississauga, Mississauga, Ontario, Canada

**Keywords:** Coexistence theory, competition, nonlinear dynamics, transients, periodic environments, seasonality, theoretical ecology

## Abstract

Despite the rapid pace of global change altering temporal environmental patterning, we lack a general understanding of how periodic environments structure ecological communities. In fluctuating environments, nonlinear dynamics associated with temporal trade-offs between competing species can create the potential for both niche differentiation (coexistence) and seemingly unexpected outcomes (exclusion) that deviate from deterministic coexistence theory. Yet, the mechanisms behind these outcomes are not fully understood. Here, we show that periodic fluctuations between times of high and low growth (e.g., seasons), and adaptive temporal trade-offs within and between species, can drive counterintuitive over- and under-performance of competing species. Most notable is the counterintuitive outcome of seasonally-mediated competitive exclusion that would not occur in either season alone, but is rather the direct result of environmental variability itself. We find that seasonal trade-offs in species’ growth rates, seasonal differences in competition strength, and functional similarity between competing species have the potential to drive nonlinear responses in coexistence to changing seasonality under global change. These biological conditions collectively influence our model’s transient dynamics, further explaining the mechanisms behind counterintuitive outcomes and highlighting the importance of non-equilibrium theory for global change ecology. Importantly, the seasonal patterns and species’ trade-offs that magnify these results are biologically realistic, therefore providing important insight into the implications for the maintenance of biodiversity under global change.

## Introduction

Periodic environmental patterns – occurring over a range of temporal scales – are ubiquitous across the globe (Vasseur and Yodzis 2004; Dillon et al. 2016; Mougi 2020). These periodicities are generally characterized by repeated fluctuations in abiotic conditions, from daily or yearly temperature and rainfall patterns to multi-year cycles characterized by periods of wet and dry conditions (e.g., ENSO). Across time scales, these abiotic periodicities drive accompanying biotic responses and can thus be characterized by periods of high growth and productivity, followed by periods of low growth and productivity. However, these environmental signals are changing due to climate change – often driving temporal asymmetries between these high- and low-growth periods. For example, northern-hemisphere winters are becoming shorter in length and more moderate (Sharma et al. 2019; Woolway et al. 2021), and weather patterns on other scales are becoming more variable and unpredictable (Yeh et al. 2009; Dillon et al. 2016). Other anthropogenic impacts are also changing abiotic periodicities, such as dam development altering seasonal water flows through entire river basins, and land modification (e.g., channelization) altering the natural waxing and waning of riverine processes (Arias et al. 2012). This makes it crucial that we develop an understanding of the role these (changing) environmental patterns play in regulating species interactions and the maintenance of biodiversity.

Environmental periodicities and associated biotic responses likely have important, yet poorly understood, implications for the structure and stability of food webs and multi-species communities (White and Hastings 2020). Specifically, variation in the structure of species interactions is likely driven by differential adaptive or behavioural responses between interacting species, with dynamical implications cascading to whole food webs (Namba 1984; Chesson et al. 2004; Mathias and Chesson 2013; McMeans et al. 2016, 2020; Scranton and Vasseur 2016). For example, competing species often have differential coping mechanisms for periods of low resource availability that are associated with trade-offs in growth potential under favourable conditions (Chesson and Huntly 1997). Simultaneously, abiotic conditions determine species’ activity rates and habitat use, which can fundamentally alter the interaction strengths (e.g., niche overlap and interspecific competition) between species through time (Chesson and Huntly 1997; Li and Chesson 2016). While research has discovered various mechanisms for species coexistence in variable environments (e.g., storage effect, relative nonlinearity, equalizing mechanisms (Armstrong and Mcgehee 1980; Amarasekare 2003; Chesson 2018; McMeans et al. 2020)), a general theory for coexistence in continuously changing environments (e.g., seasonal) is still being developed (Abrams 2022).

Indeed, recent research has suggested that periodic environmental variation (e.g., seasonality) can have a large influence on species interactions and even promote coexistence (Namba 1984; Namba and Takahashi 1993; Litchman and Klausmeier 2001; McMeans et al. 2020; Mougi 2020), while other research has shown that environmental variability can mediate a range of coexistence outcomes (Cushing 1980; Schreiber 2021; Abrams 2022; Scott et al. 2023). McMeans et al. (2020) suggest that annual fluctuations in niche partitioning between two competing predators can allow for important trade-offs that promote coexistence, and that under certain conditions, this seasonally-mediated coexistence can be highly sensitive to climate change. Recently, Scott et al. (2023) expanded this theoretical foundation to suggest fluctuation-mediated coexistence outcomes may be ubiquitous across temporal scales – also importantly highlighting the interrelated nature of relative environmental and biotic time scales (Bieg et al. 2023; Scott et al. 2023; Abbott et al. 2024). Relatedly, the theoretical literature is beginning to show that environmental variation broadly can interact with underlying nonlinearities to drive unexpected results, including fundamentally altered or novel attractors in dynamic ecological systems (Hastings et al. 2018, 2021; Bieg et al. 2022, 2023; Morozov et al. 2024). Thus, periodic environmental variation is a recipe for nonlinearities and potentially counterintuitive effects. Despite this initial theory, the potential for – and drivers of – nonlinear and/or unexpected effects on species coexistence in periodic environments is not well understood. This is an especially critical area in the face of global change altering these periodicities.

Here as a first step towards exploring the potential for counterintuitive effects of periodic environments, we expand on previous theory that found different qualitative outcomes of changing periodicities on species coexistence with temporally-variable biotic rates (McMeans et al. 2020; Scott et al. 2023). Hereafter, we refer to these fluctuations as “seasonal” for simplicity but note that our insights ought to be relevant over a range of time scales as shown by Scott et al. (2023). We focus on a particularly interesting – and counterintuitive – outcome where small changes in seasonality (e.g., modifying the relative length of seasons of high and low growth, or seasonal “symmetry”) can precipitously alter coexistence and drive exclusion even when stable coexistence ought to occur under all conditions experienced based on deterministic theory (i.e., in both seasons separately) (Scott et al. 2023). By exploring these outcomes analytically, we show that counterintuitive competitive over- and underperformance (i.e., a species does better or worse than a linear baseline) are a result of transient dynamics that are magnified by temporal trade-offs in species’ growth and competition. Specifically, seasonal growth trade-offs between species and seasonal differences in competition strength drive nonlinear responses in coexistence to changing seasonality, and this is further amplified by the functional relatedness of competing species. Importantly, these results occur more readily under trade-offs and seasonal conditions that are biologically plausible, suggesting that as climate change alters seasonal patterns, we may alarmingly expect to find counterintuitive patterns in species coexistence that our current theory is unequipped to predict.

## Methods

### Seasonal competition model

We use a seasonally-variable two species Lotka-Volterra competition model first introduced in McMeans et al. (2020) (also see Scott et al. (2023)), but loosen their biological parameter constraints for a more general theory. Specifically, we use an iterated step function of the classic Lotka-Volterra competition model (*f(t)* for two competing species *j* and *k*) to incorporate two distinct, repeating “seasons” with altered growth and competition strengths. Here, species’ densities are modelled by *f(t)* for two competing species *X*_*j*_ and *X*_*k*_ in season *s*:

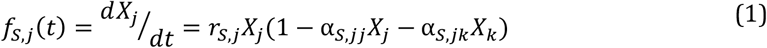

Where *r*_*s,j*_ is the intrinsic rate of population growth for species *j, a*_*s,jj*_ is the intraspecific competition rate for species *j* and *a*_*s,jk*_ is the strength of interspecific competition of species *k* on species *j*. Each of these parameters can change according to season *s*.

While we loosen McMeans et al. (2020)’s biological constraints regarding temporal changes in activity rates, resource availability, and habitat use, we assume some differentiation between species’ growth in the two periods. Therefore, we define repeated fluctuations between a high-growth (HG) and low-growth (LG) season as follows:

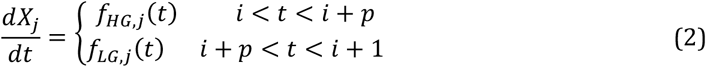

where time, *t*, runs continuously from 0 to *t*_*END*_, with two discrete “seasons” within each time unit, repeating according to *i*. In other words, seasons are discrete but dynamics within each season are continuous. Here, *i* defines the time unit, or period (e.g., year) from 0 to the length of the model run (i.e., the number of repeating periods, generally enough to reach an asymptotic state); *p* is length of the high-growth season as a proportion of each time unit (leaving (1-*p*) as the proportion of each time unit under low-growth conditions); and *j* represents either competing species *j* or *k*. We impose the condition that *r*_*HG,j*_ > *r*_*LG,j*_, for both species, to represent the HG (high-growth) and LG (low-growth) seasonal patterning. Using this model, we can vary the parameter *p* as also done in McMeans et al. (2020) and Scott et al. (2023) to determine the effect of changing periodicities on species coexistence.

Numerical simulations are coupled such that species’ densities at the end of each season become initial values for the next season, at which point the parameters mentioned above change instantaneously. After a sufficiently long transient period, all numerical simulations settle at an asymptotic state, such that species’ densities fluctuate within each time unit (e.g., year) around a mean that remains constant. All analyses were done using Wolfram Mathematica (version 12.3.1.0) and numerical simulations were evaluated using the built-in ODE solver “NDSolve.”

### Approximate analytical solution for seasonal competition model

Scott et al. (2023) developed a simple analytical approximation of the seasonal model’s asymptotic dynamics (also see Litchman and Klausmeier (2001) for a similar approximation under fast environmental forcing; see Appendix A1). In a simplified sense, the coupled vector fields of high- and low-growth seasons allow for a time-weighted “average” of linearized dynamics throughout state space. In the limit of fast forcing (rapid switching between discrete “seasons”, i.e., vector fields), this linearized approximation necessarily converges to the true dynamics (Hadeler and Hillen 2007), but Scott et al. (2023) show that this approximation replicates numerical solutions remarkably well across a range of time scales. Here, we harness this analytical solution to determine biological mechanisms underlying counterintuitive competitive performance, along with numerical simulations for confirmation.

This approximation gives us coexistence criteria resembling the original Lotka-Volterra criteria for a seasonally-variable competitive interaction (see Table 1 for a subset of outcomes and Table A1 for the full set of criteria). Notably, these criteria highlight that coexistence is dependent on seasonal-growth-scaled inter-versus intra-specific competition strength (whereas the classical Lotka-Volterra criteria are simply dependent on inter-versus intra-specific competition strength; that is, coexistence occurs when *a*_*kj*_ /*a*_*jj*_ < 1 for both species). Here, we use these criteria in what follows to explore seasonally-mediated changes in coexistence (i.e., bifurcations driven by *p*; for an example of a bifurcation using these isoclines see Figure A1).

**Table 1.**
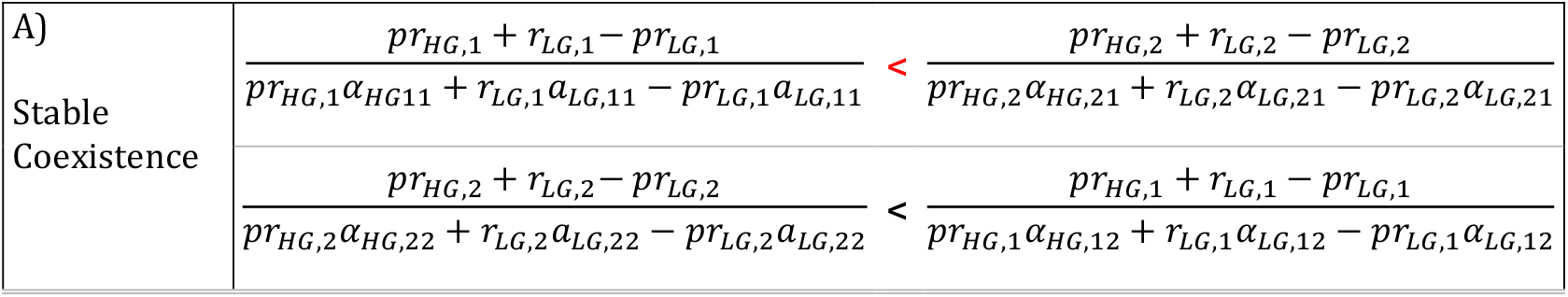
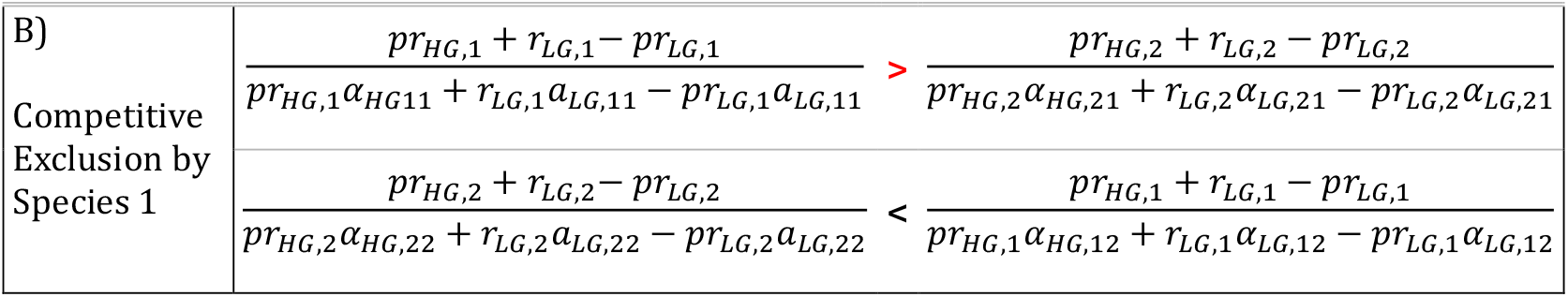
Coexistence criteria for seasonal environments from approximated isocline solutions (Appendix A1) as an extension to classical L-V competition theory, highlighting the difference between (A) stable coexistence and (B) competitive exclusion of species 2 by species 1. In the case of seasonally-mediated competitive exclusion (as shown in Figure 1), we find the scenario where both seasons individually (i.e., boundary conditions at *p*=0 or 1) satisfy (A) but intermediate *p* satisfies (B), such that changing *p* drives a flip in the inequality highlighted in red.

|  |  |
| --- | --- |
| A)<br>Stable<br>Coexistence | $\frac{pr_{HG,1} + r_{LG,1} - pr_{LG,1}}{pr_{HG,1}\alpha_{HG,11} + r_{LG,1}\alpha_{LG,11} - pr_{LG,1}\alpha_{LG,11}} < \frac{pr_{HG,2} + r_{LG,2} - pr_{LG,2}}{pr_{HG,2}\alpha_{HG,21} + r_{LG,2}\alpha_{LG,21} - pr_{LG,2}\alpha_{LG,21}}$ |
| | $\frac{pr_{HG,2} + r_{LG,2} - pr_{LG,2}}{pr_{HG,2}\alpha_{HG,22} + r_{LG,2}\alpha_{LG,22} - pr_{LG,2}\alpha_{LG,22}} < \frac{pr_{HG,1} + r_{LG,1} - pr_{LG,1}}{pr_{HG,1}\alpha_{HG,12} + r_{LG,1}\alpha_{LG,12} - pr_{LG,1}\alpha_{LG,12}}$ |

|  |  |  |
| --- | --- | --- |
| B)<br>Competitive<br>Exclusion by<br>Species 1 | $\frac{pr_{HG,1} + r_{LG,1} - pr_{LG,1}}{pr_{HG,1}\alpha_{HG,11} + r_{LG,1}\alpha_{LG,11} - pr_{LG,1}\alpha_{LG,11}} > \frac{pr_{HG,2} + r_{LG,2} - pr_{LG,2}}{pr_{HG,2}\alpha_{HG,21} + r_{LG,2}\alpha_{LG,21} - pr_{LG,2}\alpha_{LG,21}}$ | |
| | $\frac{pr_{HG,2} + r_{LG,2} - pr_{LG,2}}{pr_{HG,2}\alpha_{HG,22} + r_{LG,2}\alpha_{LG,22} - pr_{LG,2}\alpha_{LG,22}} < \frac{pr_{HG,1} + r_{LG,1} - pr_{LG,1}}{pr_{HG,1}\alpha_{HG,12} + r_{LG,1}\alpha_{LG,12} - pr_{LG,1}\alpha_{LG,12}}$ | |

### Understanding the effects of changing seasonality

This periodic model composed of two discrete seasons (e.g., low- and high-growth seasons, with continuous dynamics within each season) repeated over time (i.e., years), allows us to make predictions regarding the effect of changing seasonality based on the extreme endpoints (i.e., conditions unique to each season). The endpoints (i.e., *p* = 0 or 1) are the environmental and biological conditions characteristic of each season (low-growth, LG, and high-growth, HG) and represent classical Lotka-Volterra coexistence outcomes based on those conditions for static environments. Between these endpoints, 0 < *p* < 1 represents fluctuating environments with different patterning (where *p* is the proportion of each time step that is under HG conditions). As such, we can think of changing seasonality (or, more generally, temporal patterning) via *p* (also see McMeans et al. 2020 and Scott et al. 2023) as moving from one extreme (each year = 100% LG conditions) to the other (each year = 100% HG conditions), and logically predict a **linear** path between these as a reasonable first approximation (i.e., as an averaging of the seasonal conditions) and will serve as a baseline which we can compare nonlinear outcomes to.

We can use the aforementioned analytical approximation to evaluate the effects of changing seasonality and determine when and why they differ from this linear baseline. Scott et al. (2023)’s analytical approximation allows us to follow the continuous effect of changing *p*, when the attractor does not track a linear prediction between two endpoints or boundary conditions at *p*=0 and *p*=1. In cases where *p* itself drives bifurcations in our seasonal model (e.g., seasonally-mediated coexistence outcomes; most often a transcritical bifurcation as an interior solution enters or exits positive state space), we can then use this approximation to track the asymptotic behaviour through parameter space in a way that is analytically tractable. This allows for a continuous measurement of potential nonlinear changes in the “attractor” even when this attractor is in negative state space and numerical solutions would fall to an axial solution. Therefore, in what follows, we occasionally follow negative densities, although biologically impossible, to quantify the extent of the nonlinearity imposed by biological conditions, relative to the linear path between *p*=0 and *p*=1 endpoints.

### The problem: Nonlinear responses and competitive underperformance

Each panel in Figure 1 displays two contrasting changes in the asymptotic state (mean densities of two competing species) in response to changing season lengths (changing *p*): 1) the linear prediction between deterministic endpoints (grey lines), and 2) a nonlinear change in species’ densities deviating from this trajectory (black lines). Here, each endpoint yields stable coexistence and we consider how changing *p* changes the competitive outcome, relative to these linear predictions. Figure 1A and B shows two strong nonlinear responses to changing season length, that deviate away from the linear predictions in counterintuitive ways. Here, we see one case that drives species 2 to low densities (Figure 1A), far past what would be expected based on either endpoint, and another that drives species 2 to local extinction over a range of intermediate *p* (Figure 1B). The directional orange arrows in Figure 1 stylistically represent this over- and under-performance of each species – that is, the extent of deviation away from the linear prediction.

**Figure 1.**
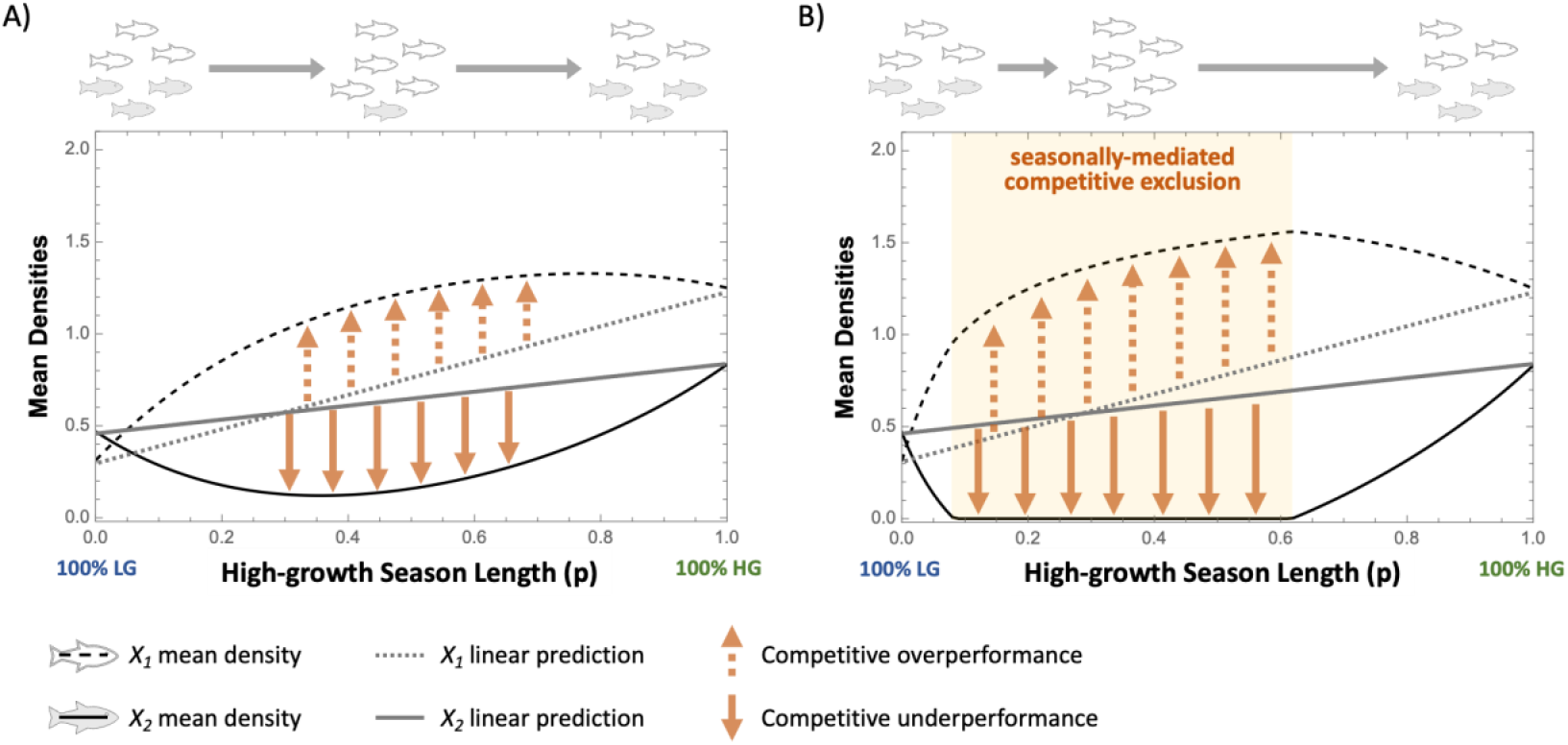
Counterintuitive effects of changing periodic signals (high-growth season length as a proportion of each time unit, *p*), relative to linear predictions (grey lines) based on the endpoints/boundary conditions. Boundary conditions in both cases define stable coexistence at the endpoints (no seasonality) but this is altered at intermediate season lengths such that competing species can have both positive and negative responses to the same changing condition (i.e., “competitive over- /under-performance” indicated by orange arrows). A) nonlinear changes in density with altered seasonality; B) seasonally-mediated competitive exclusion (an extreme case of (A) driven through the X_2_=0 boundary). Here, even though both boundary conditions allow for stable coexistence (parameters in each season facilitate coexistence between both species), fluctuations between these seasonal conditions drives one species (species 2) to exclusion.

Importantly, the outcomes shown in Figure 1 suggest that temporal variation can drive unexpected (from an endpoint equilibrium perspective) results and potentially drive fundamentally different coexistence outcomes compared to static environments. Furthermore, if we imagine the case where a species is better adapted to at least one of the seasons, then indeed these curves also have the biologically counterintuitive result that this same species decreases in some situations (i.e., changing *p* over some range) even when there is an increase in the season it is better adapted to. For example, increasing *p* from 0.6 onwards in Figure 1 causes the LG-favoured species (species 2) to increase while the HG-adapted species (species 1) decreases. Since climate change is threatening to alter the nature of common periodicities (e.g., changing seasonal durations, often with high-growth seasons lengthening and low-growth seasons decreasing), we are motivated to understand what governs the strength of this nonlinearity and when we might see these remarkably counterintuitive competitive outcomes in a changing climate. In what follows, we explore biological conditions that ought to amplify these counterintuitive results. We also explore how these conditions dictate transient dynamics and their role in driving counterintuitive, nonlinear coexistence outcomes in fluctuating environments.

### Towards biological rules for seasonally underperforming competitors

Towards understanding biological conditions that may amplify the counterintuitive patterns displayed in Figure 1, we can deduce simple mathematical rules using our analytical coexistence conditions (Table 1, Appendix A1) along with numerical simulations that track this competitive over/underperformance. The nonlinear outcome we are interested in here necessarily occurs when the inequalities defining the coexistence condition (Table 1A) fundamentally change to those of the exclusion condition (Table 1B) as *p* increases from 0, and then again satisfy the coexistence condition at *p* = 1. That is, the condition at the top of Table 1, where the left-hand-side (LHS) is less than the right-hand-side (RHS), is satisfied at *p* = 0 and 1, but LHS > RHS at intermediate *p*. To identify mechanisms underlying these counterintuitive outcomes, we explore conditions enabling this result based on the relative differences in changes to LHS and RHS when increasing *p* from 0 – i.e., towards a flip in the inequalities. Immediately, we can deduce several insights into what ought to drive this relationship between the LHS and RHS terms in Table 1 (these are stated more formally in Appendix A2).

First, despite having high- and low-growth seasons defining general growth patterns over time, we see that seasonal asymmetries or trade-offs between species’ *relative* growth rates can drive this nonlinear effect. With increasing *p*, the numerator ought to increase the LHS more than the RHS when the growth rates are asymmetric such that *r*_*HG,1*_ > *r*_*HG,2*_ and *r*_*LG,1*_ < *r*_*LG,2*_. These conditions thus make it more likely to see the inequality change in the coexistence conditions (Table 1) with increasing *p* from 0. We refer to this as the **seasonal growth trade-off effect**. In other words, competing species being *relatively* better performers in different seasons may be a factor leading to counterintuitive coexistence outcomes in periodic environments.

Second, we see that strong competition during the low-growth season may also drive this effect. Given our boundary conditions (i.e., *α*_11_ *> α*_21_ in both seasons, meaning the denominator for the LHS ought to be more affected by changing *p* than that of the RHS), we ought to see a more rapidly increasing LHS when *p* increases from 0 when *α*_*HG*,11_ < *α*_*LG*,11_. We refer to this result as the **seasonal competition effect**. That is, differences in competition strength between seasons can lead to counterintuitive competitive outcomes, despite the ratio of inter-to intra-specific competition strength always predicting stable coexistence based on classic coexistence theory criteria.

Finally, we note that the baseline coexistence criteria (*α*_11_*/α*_21_) may alter the magnitude of the nonlinear effect when the ratio is closer to 1 (i.e., baseline, or within season, inter- and intra-specific competition strengths are more similar). The RHS denominator in Table 1 can be made more likely to allow for the required flip in the coexistence condition if the interspecific competition strengths (*α*_21_, for either season) in the RHS denominator are nearly equal to the intraspecific competition strengths (*α*_11_; note, interspecific competition strength must remain < *α*_21_ to retain the boundary conditions). In this case, the RHS denominator becomes larger and more similar to the necessarily larger LHS denominator. This would then be expected to amplify the effect of the asymmetries above – particularly those of the growth rates. We refer to this amplifying effect as the **“sister species” effect** as similar species would be expected to have more similar intraspecific and interspecific competition effects.

Below we use this logic to explore these nonlinear responses numerically. We quantified the counterintuitive effect size using our analytical approximation (Appendix A1) by calculating the maximum deviation of species 2’s (the species that is prone to exclusion) “equilibrium” (i.e., the mean density at the asymptotic state) from a linear baseline based on the seasonal endpoints, across values of *p*. This underperformance can be seen by the orange arrows in Figure 1, though here we track the approximation’s equilibrium into negative state space (if it crosses this boundary; i.e., through a transcritical bifurcation) for a continuous measurement of counterintuitive effect size. Effectively, this determines the magnitude of species 2’s competitive underperformance, given the linear baseline between boundary conditions such that a larger effect size represents an increased potential for species 2 to become locally extinct at intermediate *p*.

### Seasonal Growth Trade-offs

First, our results confirm that seasonal growth rate differentials amplify the nonlinear effect of changing *p*, referred to above as the **seasonal growth trade-off effect** (Figure 2). Figure 2 shows that increasing inter-specific growth rate differentials in each season (i.e., increasing *r*_*HG,1*_ and *r*_*LG,2*_ based on conditions identified above and in Appendix A2), while holding other rates constant, amplifies the nonlinear effect size. Note that the inflection point does not perfectly align with the other species’ growth rate due to underlying seasonal differences in competition strength, which are held constant but necessary for some seasonal differentiation in the underlying attractors. This growth rate effect is also controlled by underlying inter- and intra-specific competition strength differences, such that the effect of growth rate differentials is amplified through the **“sister species” effect** described above.

**Figure 2.**
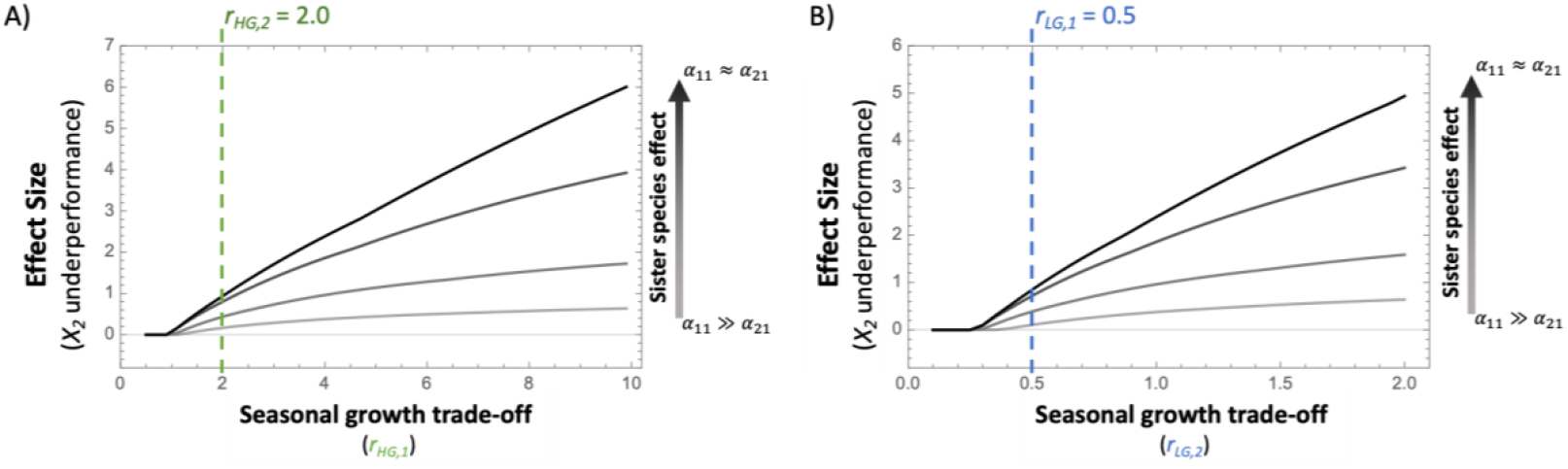
Interspecific *r*-asymmetries (i.e., seasonal growth trade-offs) amplify counterintuitive competitive performance, the magnitude of which depends on boundary conditions (defined by seasonal inter-vs. intra-specific competition strengths). Specifically, increasing the inequalities *r*_*HG,1*_ *> r*_*HG,2*_ and *r*_*LG,2*_ > *r*_*LG,1*,_ increases the magnitude of the counterintuitive effect across changing seasonal proportions. The effect size is measured as the maximum difference across *p* = (0, 1) between species 2’s mean density and the linearly predicted equilibrium based on boundary conditions (i.e., the maximum underperformance). Note that *r*_*HG*_ > *r*_*LG*_ for both species, and in both A) and B), ***α***_***LG***_ > ***α***_***HG***_. The sister species effect (when ***α***_**21**_/***α***_**11**_ ≈ 1) amplifies the counterintuitive effect of *r*-asymmetries even further, shown by grey-to-black shading of lines (changing ***α***_**21**_/***α***_**11**_ ratio from 0.45 to 0.9).

Intriguingly, this result suggests that even when species 2 has a relative growth advantage over species 1 in the low-growth season, and species 2 would be expected to persist under all conditions, it may be pushed towards local extinction even in situations where most of the time is spent in those relatively favourable conditions (low-growth season). Similarly, even when at a relative growth disadvantage in high-growth periods, the density of species 2 may actually increase with further increases in high-growth seasonal duration (as *p* approaches 1) while species 1 – with the growth advantage in the high-growth periods – counterintuitively declines (Figure 1).

### Seasonal Differentiation in Competition Strengths

Next, we see that seasonal differences in competition strength can indeed also drive this counterintuitive response, referred to here as the **seasonal competition effect**. In Figure 3, we altered the competition strengths in each season, such that interspecific competition (*α*_21_) was held as a constant proportion of species 1’s intraspecific competition strength (*α*_11_; i.e., controlling for the sister species effect and boundary condition *α*_21_/*α*_11_). We see that when increasing inter- and intra-specific competition strengths together, the nonlinear effect of changing *p* is amplified by strong seasonal differences in competition (Figure 3). Specifically, weak competition in the high-growth season (Figure 3A) and strong competition in the low-growth season (Figure 3B) drive underperformance of species 2. Additionally, when *α*_21_/*α*_11_ is held closer to 1 (i.e., *α*_11_ ≈ *α*_21_), the sister species effect again amplifies the effect of changing competition strength – particularly when LG competition is strong (Figure 3).

**Figure 3.**
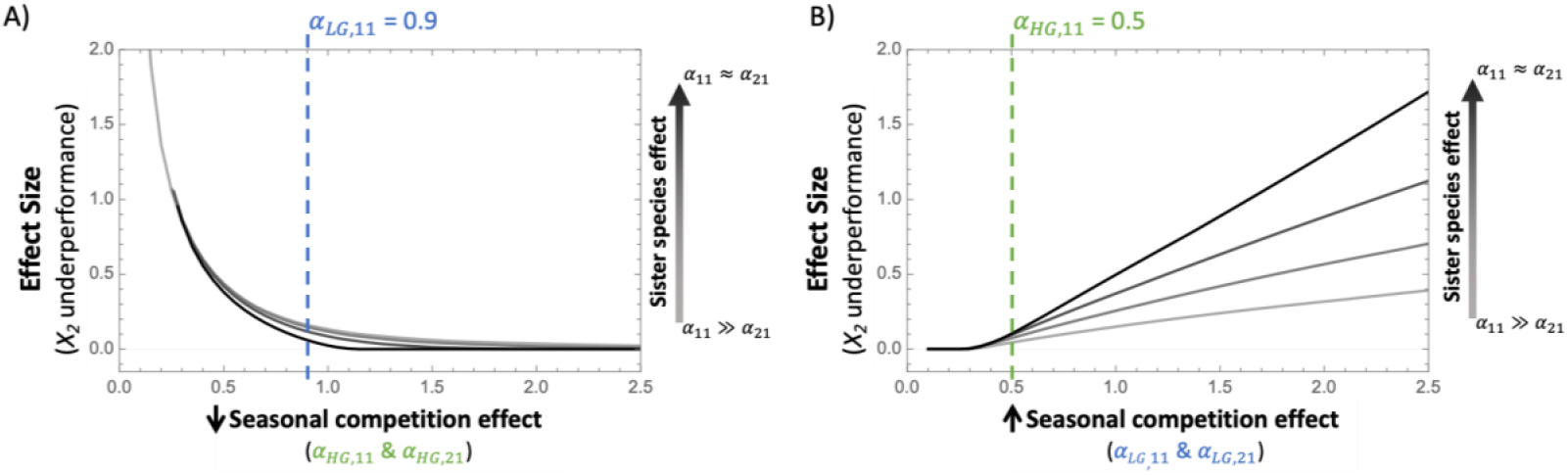
Effect of changing seasonal competition differences between high and low growth seasons (varying ***α***_S,11_, with ***α***_S,21_ held at constant proportions) on the magnitude of the counterintuitive effect across changing seasonal proportions. The effect size is measured as the maximum difference across *p* = (0, 1) between species 2’s mean density and the linear prediction. ***α***_***LG***_ > ***α***_***HG***_ magnifies the underperformance of species 2, given modest seasonal growth rate trade-offs. The sister-species effect (when ***α***_**21**_/***α***_**11**_ ≈ 1) amplifies the nonlinear effect of competition differences even further, shown by grey-to-black shading of lines (changing ***α***_**21**_/***α***_**11**_ ratio from 0.45 to 0.9).

This result implies that large seasonal differences in competition strength – even when inter-versus intra-specific competition strength in both seasons ought to always facilitate stable coexistence – may drive nonlinear effects of changing seasonality. Specifically, this occurs when competition (both inter- and intraspecific) is strong in low-growth seasons, and relatively weak under high-growth conditions. Again, the ratio of inter-versus intra-specific competition strength may dampen (Figure A2.A,B) or strengthen (Figure A2.C,D) this pattern based on the sister species effect.

### Unpacking the dynamics behind competitive underperformance

We have identified general relationships between species’ seasonal growth trade-offs and coinciding seasonal patterns in competition strength that ought to amplify nonlinear effects of changing seasonality. Our results suggest the plausible notion that environmental conditions causing inversely related growth rates and competition strengths across seasons (i.e., low growth and strong competition, high growth and weak competition) may be especially prone to nonlinear competitive outcomes. Importantly, this suggests that seasonal asymmetries in biological rates may trigger unexpected population declines and potentially lead to local extinction. Next, to further unpack why we see these counterintuitive outcomes and understand these asymmetries collectively, we can step away from our boundary conditions and view these results in the phaseplane (Figure 4). We show that the nonlinear changes in the attractor with changing *p* are due to the coupling of transient dynamics under each season (i.e., the underlying vector fields), and explore how asymmetries in species’ growth and competition trade-offs act together to regulate these dynamics.

**Figure 4.**
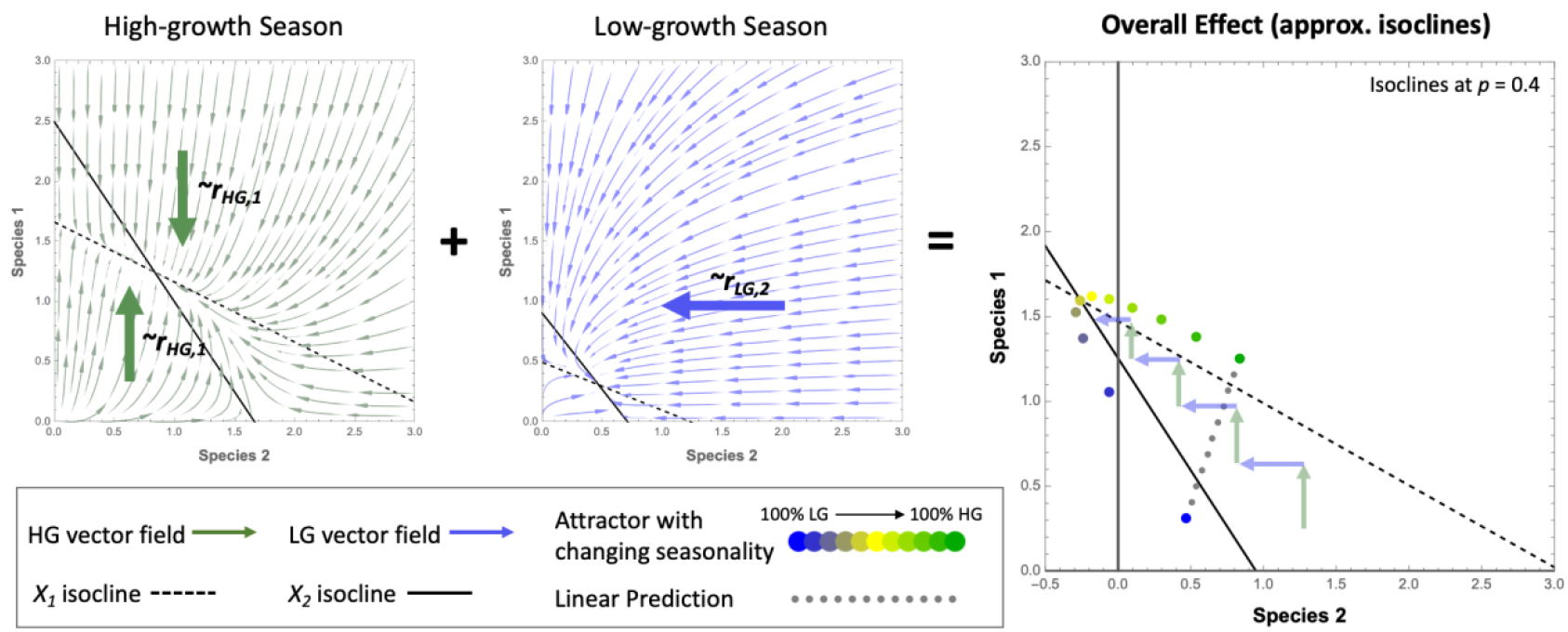
Seasonal isocline geometry and underlying vector fields, with directionality and magnitude of the vector fields proportional to growth rates. Together, these vector fields act like a ratchet to drive the overall attractor towards the boundary (i.e., a “net effect” that is overall negative in the *X*_*2*_ direction). Temporal growth asymmetries “linearize” the seasonal vector fields and amplify a “ratchet effect”. The rightmost panel shows the nonlinear path in the attractor (coloured dots) with changing *p* (for reference, the linear prediction between boundary conditions is indicated by gray dots) leading to seasonally-mediated competitive exclusion at intermediate *p*, as a result of this ratchet effect.

Figure 4 shows the seasonal (boundary) conditions in the phaseplane, including underlying vector fields. The directionality of the vector fields are governed by growth rates, whereas the isocline geometry and equilibrium densities are only dependent on inter- and intra-specific competition strengths. In a sense, high *r*_*HG,1*_ “linearizes” the vector field in the vertical direction (i.e., parallel to the species 1 axis) in the high-growth season and high *r*_*LG,2*_ “linearizes” the low-growth season’s vectors in the horizontal, species 2 direction. The strength of these vectors are roughly proportional to the growth rates, regardless of the competition strengths (though the competition rates clearly alter the directionality of the vector fields near the equilibria). Given enough seasonal differentiation in the competitive interaction to separate the seasonal equilibria in the phaseplane (i.e., increase the “room” for transient dynamics to generate nonlinearities), these growth rate differentials create seasonal asymmetries in the species’ numerical responses and push the system away from either equilibrium. Collectively, this drives a “ratchet effect” away from the predicted equilibrium (i.e., the linear baseline) by coupling the seasonal vectors over time (also see Greyson-Gaito et al. (2023) for a related result in consumer-resource interactions). This ratchet pushes the attractor to low species 2 densities, until the point at which within-season dynamics negate each other and the system reaches an asymptotic state. Figure 4 shows a qualitative representation of this ratchet working to dictate the overall trajectory in our seasonal model. Essentially, these “linearized” seasonal vector fields couple together to create nonlinearities in the overall vector field under seasonal variation and suppress species 2.

In the example shown in Figure 4, both inter- and intraspecific competition are stronger in the low-growth season than the high-growth season (i.e., *α*_*LG*_ > *α*_*HG*_ as noted as one of our conditions above) and thus the low-growth season’s equilibrium densities are suppressed towards the origin relative to the high-growth season’s equilibrium. However, densities in the seasonal model do not simply fluctuate directly between the two equilibria. The geometrical configuration shown here makes it so that the high *r*_*LG,2*_ effectively speeds up the “negative growth” in the low-growth season (hence the arrow direction in Figure 4) and ensures this detrimental (for species 2) “ratchet effect.” It is also worth noting that if *α*_11_ ≈ *α*_21_ then the isocline intercepts on the species 1 axis (y axis) in Figure 4, for both seasons, would be closer together, moving the seasonal equilibria closer to this axis. This would further increase the likelihood of species 2 getting pushed through the 0-boundary due to any nonlinearities in the attractor’s response, especially with the “negative” vectors during the low-growth season. Thus, we can visualize how environmental fluctuations between seasons with biological trade-offs can drive unexpected nonlinear coexistence outcomes based on how these conditions interact with each other.

Another way to biologically interpret the ratchet effect displayed here is that large numerical responses in species 1 during the high-growth seasons (i.e., the green arrows in Figure 4), resulting in increased species 1 densities leading into the low-growth periods, subsequently lead to large numerical responses in the low-growth seasons – specifically large, negative effects on species 2 (i.e., blue arrows in Figure 4) due to the multiplicative effect of density and competitive effect (Equation 1). While true even under moderate inter-specific competition strength (*α*_*LG*,21_), *α*_*LG*,21_ ≈ *α*_*LG*,11_ clearly further magnifies this effect in the low-growth seasons. Here, strong growth responses by species 1 (i.e., *r*_*HG,1*_ >> *r*_*HG,2*_) during the high-growth seasons further amplify this ratchet effect. That is, a strong seasonal growth trade-off effect feeds into a strong seasonal competition effect, generating a positive feedback of competitive over- and under-performance.

## Discussion

We have identified scenarios where changing environmental periodicities can drive nonlinear responses in species’ densities and result in counterintuitive competitive outcomes (Figure 1). These outcomes have clear implications for the maintenance of biodiversity under global change. Specifically, we have identified biological conditions whereby certain species can competitively underperform and be pushed near or to local extinction, even in environments that deterministically (i.e., without environmental fluctuations) ought to facilitate coexistence at all times (over the full range of environmental conditions experienced within a year). Alarmingly, we note that the conditions driving these nonlinear responses may be biologically realistic.

Seasonal trade-offs in growth (Figure 2) and competition (Figure 3) can alter the performance of competing species in counterintuitive ways. The trade-offs and asymmetries identified here may all act together to drive counterintuitive over- or underperformance of competing species in changing environments. Specifically, we refer to the effects of these trade-offs as: **seasonal growth trade-off effect, seasonal competition effect**, and **sister species effect** (Table A2). Under these biotic conditions, competing species with differential growth responses to environmental variation (i.e., each has *relatively* better performance or fitness in different seasons, despite there being high- and low-growth seasons) and differential competition strengths between seasons (especially if competition is stronger in the low growth season) may have fundamentally altered coexistence outcomes than would be expected in a static environment. We note that while not all of these biological conditions are simultaneously necessary to drive the results seen here, some seasonal and inter-specific differentiation is clearly needed to allow for a nonlinear response in competitive outcomes (i.e., the attractor’s path shown in Figure 4). Here, we have also begun to unpack the dynamical mechanisms behind these counterintuitive results and highlight that they are driven by interactive transient dynamics that couple together to create a “ratchet effect” on the resulting attractor (Figure 4).

Interestingly, the trade-offs between competing species that we have identified seem biologically likely. Species commonly have growth or performance trade-offs between different environmental conditions (e.g., periods of changing temperature or resource availability) which can equalize their competitive abilities (Armstrong and Mcgehee 1980; Chesson and Huntly 1997; Chesson 2018; McMeans et al. 2020). McMeans et al. (2020) synthesized examples of several fish competitors that show temporal differentiation in growth, activity, niche overlap and competitive ability, with trade-offs within species pairs associated with their thermal guilds (lake trout and smallmouth bass, burbot and lake trout, Arctic charr and brown trout, European perch and roach). Furthermore, plasticity in the feeding strategies of neotropical fish has been shown to allow for niche divergence during periods of high resource availability (Neves et al. 2021), which would lower interspecific (through less niche overlap) and intraspecific (from increased resource availability) competition during these times. This suggests that, depending on the overlap and directionality of these various temporal trade-offs, these species may display biological conditions in line with our enhanced nonlinearity criteria. These general biological conditions are not restricted to aquatic ectotherms, though. For example, coexistence in desert rodent communities is supported by temporal trade-offs in foraging efficiency (Brown 1989), and it is common for competitors with similar niches to have differential growth strategies or other forms of temporal differentiation (Chesson 2018). Together these examples describe well-known coexistence mechanisms (e.g., equalizing and stabilizing mechanisms), but our results suggest that in some cases these biological conditions (e.g., trade-offs) may actually have unexpected, detrimental effects for coexistence.

Trade-offs can be associated with differential competitive abilities between seasons, however species have also adapted behavioural adaptations to cope with periods of harsh conditions or low resource availability in different ways (Chesson and Huntly 1997). These behavioural strategies may act to reinforce the nonlinear responses we see here (e.g., increased niche overlap or range contraction that would increase competition during harsh periods) (McMeans et al. 2020), or alternatively counteract a species’ potential for competitive underperformance (e.g., torpor/hibernation or migration that drives niche divergence when competition would otherwise be strong) (Schoener 1982; Cáceres 1997; Harabiš et al. 2012). Depending on how these adaptations alter niche overlap between species, particularly during sub-optimal times, they may act as stabilizing mechanisms to counteract competitive underperformance in periodic environments. The implications of global change – with associated alterations in environmental periodicities – for these competitive trade-offs remain unclear, however we suggest that competing species with growth trade-offs that equalize them and with similar niches (i.e., without stabilizing mechanisms that allow for niche divergence), may be most susceptible to these nonlinear effects.

These nonlinear responses to changing environmental periodicities, and realistic biological conditions that can magnify them, have clear implications for global change. Notably, changing durations of annual seasons (e.g., cold or dry seasons) as well as temporal patterning on other time scales (i.e., times of low/high productivity or growth from daily to decadal periods; Scott et al. 2023) may have drastic effects on coexistence and community structure. Indeed, this work shows that the conditions necessary for driving counterintuitive competitive performance are related to relative differences in key traits between seasons and species and are likely to be important across temporal scales. This suggests that changing periodicities may precipitously drive species towards local extinction when they would otherwise be expected to persist – or even thrive – under all conditions. Similarly, we see that species with seasonal growth advantages can counterintuitively decline with increasing duration of their favourable season, and species with a growth disadvantage under those same conditions may increase.

In the face of rapid global change altering environmental periodicities and stretching species’ adaptive capacities to their limits, understanding these biotic trade-offs and resulting implications for the maintenance of biodiversity is critical. Between population range shifts, novel species introductions and changing biological rates, environmental periodicities and resource availability, global change may fundamentally alter competitive interactions and rewire whole communities (Penk et al. 2016; Lancaster et al. 2017; Bartley et al. 2019). Ultimately, species coexistence may be susceptible to changing environmental conditions and indeed we show here that this sensitivity can also drive highly unexpected outcomes under global change. Our ability to forecast the impacts of global change requires us to understand the mechanisms underlying potential nonlinearities and deviations from what deterministic theory predicts.

## Funding Statement

This project was supported through a Canada First Research Excellence Fund project “Food from Thought” and NSERC Discovery Grant (#400353) to KSM, and CB was supported by NSERC CGS-D funding.

## Author Contributions

All authors contributed to the development of ideas, CB, AS and KSM designed the theoretical model and analytical solution, and CB created the figures. CB wrote the initial draft of the manuscript and all authors contributed to editing subsequent revisions.

## Code Availability

Mathematica code used for analysis is available on GitHub:

## Appendix: Supplementary material for Bieg et al

*Appendix A1 is information summarized from Scott et al. (2023)*.

### A1. Isocline Approximation for coexistence criteria in periodic environments

Although the nonequilibrium model used here is mathematically intractable, we can approximate isocline solutions using some simple assumptions of linearity for an analytical approach that maps surprisingly well to numerical solutions across a range of time scales of environmental periodicities (Scott et al. 2023). Specifically, if we assume that the seasonal trajectory of our system, given some initial value, is linear and scaled by the duration of that season, then we can assume the change of a given species *j* over the high-growth season can be approximated by:

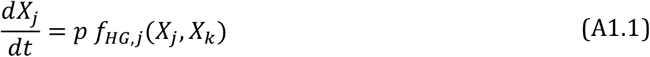

and similarly, the low-growth trajectory scales linearly as:

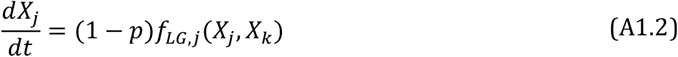

Under this assumption of linearity, we can calculate approximate solutions for the isoclines when these seasonal trajectories are equal and opposite of each other and thus the NET change in species densities over a unit of time (e.g., 1 year) ought to be zero. That is:

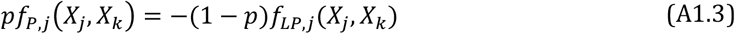

Adding in the Lotka-Volterra competition equations, *f(t)*, for two species, species 1 (*X*_*1*_) and species 2 (*X*_*2*_), and solving as functions for *X*_*1*_, we are left with the following isocline equations:

Species 1 isocline:

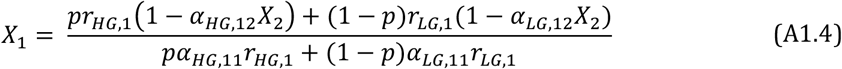

Species 2 isocline:

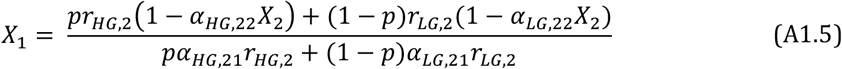

Using Equations A1.4 and A1.5 we can solve for the equilibrium solutions (i.e., A1.4 = A1.5). The nontrivial equilibrium solution is not easily comprehendible in symbolic form, but we note that it tracks the attractor with changing *p* well across a range of parameters and time-scales (see Scott et al. 2023 for a more in-depth explanation). Regardless of this, we can use the isocline solutions to develop general coexistence criteria for our periodically forced model, that geometrically and conceptually map to the original coexistence criteria for the Lotka-Volterra competition model (i.e., dependent on inter-versus intraspecific competition strengths). In our case, the isocline geometry matches that of the original Lotka-Volterra geometry and we can thus use similar rules: specifically, the relationship between seasonal-growth-scaled inter-versus intra-specific competition (Table A1).

**Table A1.** Coexistence criteria for competitive interactions in periodic environments.

|  |  |
| --- | --- |
| Coexistence | $\frac{pr_{HG,1} + r_{LG,1} - pr_{LG,1}}{pr_{HG,1}\alpha_{HG,11} + r_{LG,1}\alpha_{LG,11} - pr_{LG,1}\alpha_{LG,11}} < \frac{pr_{HG,2} + r_{LG,2} - pr_{LG,2}}{pr_{HG,2}\alpha_{HG,21} + r_{LG,2}\alpha_{LG,21} - pr_{LG,2}\alpha_{LG,21}}$ |
| | $\frac{pr_{HG,2} + r_{LG,2} - pr_{LG,2}}{pr_{HG,2}\alpha_{HG,22} + r_{LG,2}\alpha_{LG,22} - pr_{LG,2}\alpha_{LG,22}} < \frac{pr_{HG,1} + r_{LG,1} - pr_{LG,1}}{pr_{HG,1}\alpha_{HG,12} + r_{LG,1}\alpha_{LG,12} - pr_{LG,1}\alpha_{LG,12}}$ |
| Competitive<br>Exclusion by<br>Species 1 | $\frac{pr_{HG,1} + r_{LG,1} - pr_{LG,1}}{pr_{HG,1}\alpha_{HG,11} + r_{LG,1}\alpha_{LG,11} - pr_{LG,1}\alpha_{LG,11}} > \frac{pr_{HG,2} + r_{LG,2} - pr_{LG,2}}{pr_{HG,2}\alpha_{HG,21} + r_{LG,2}\alpha_{LG,21} - pr_{LG,2}\alpha_{LG,21}}$ |
| | $\frac{pr_{HG,2} + r_{LG,2} - pr_{LG,2}}{pr_{HG,2}\alpha_{HG,22} + r_{LG,2}\alpha_{LG,22} - pr_{LG,2}\alpha_{LG,22}} < \frac{pr_{HG,1} + r_{LG,1} - pr_{LG,1}}{pr_{HG,1}\alpha_{HG,12} + r_{LG,1}\alpha_{LG,12} - pr_{LG,1}\alpha_{LG,12}}$ |
| Competitive<br>Exclusion by<br>Species 2 | $\frac{pr_{HG,1} + r_{LG,1} - pr_{LG,1}}{pr_{HG,1}\alpha_{HG,11} + r_{LG,1}\alpha_{LG,11} - pr_{LG,1}\alpha_{LG,11}} < \frac{pr_{HG,2} + r_{LG,2} - pr_{LG,2}}{pr_{HG,2}\alpha_{HG,21} + r_{LG,2}\alpha_{LG,21} - pr_{LG,2}\alpha_{LG,21}}$ |
| | $\frac{pr_{HG,2} + r_{LG,2} - pr_{LG,2}}{pr_{HG,2}\alpha_{HG,22} + r_{LG,2}\alpha_{LG,22} - pr_{LG,2}\alpha_{LG,22}} > \frac{pr_{HG,1} + r_{LG,1} - pr_{LG,1}}{pr_{HG,1}\alpha_{HG,12} + r_{LG,1}\alpha_{LG,12} - pr_{LG,1}\alpha_{LG,12}}$ |
| Contingent<br>Coexistence | $\frac{pr_{HG,1} + r_{LG,1} - pr_{LG,1}}{pr_{HG,1}\alpha_{HG,11} + r_{LG,1}\alpha_{LG,11} - pr_{LG,1}\alpha_{LG,11}} > \frac{pr_{HG,2} + r_{LG,2} - pr_{LG,2}}{pr_{HG,2}\alpha_{HG,21} + r_{LG,2}\alpha_{LG,21} - pr_{LG,2}\alpha_{LG,21}}$ |
| | $\frac{pr_{HG,2} + r_{LG,2} - pr_{LG,2}}{pr_{HG,2}\alpha_{HG,22} + r_{LG,2}\alpha_{LG,22} - pr_{LG,2}\alpha_{LG,22}} > \frac{pr_{HG,1} + r_{LG,1} - pr_{LG,1}}{pr_{HG,1}\alpha_{HG,12} + r_{LG,1}\alpha_{LG,12} - pr_{LG,1}\alpha_{LG,12}}$ |

It is worth noting that the simple isocline approximation can estimate these patterns of changing *p* (e.g., seasonally-mediated changes in density or bifurcations) surprisingly well, but the geometry alone doesn’t fully explain *why* we see certain results (i.e., counterintuitive or nonlinear responses). In fact, due to the assumption of linearization, it is surprising that the approximation does work so well given nonlinear vector fields. The proof of the efficacy of this approximation can be seen in Scott et al. (2023).

### A2. Identifying biological rules for seasonally-mediated competitive exclusion

We are interested in understanding how coexistence under static low-growth conditions (*p*=0) and coexistence under static high-growth conditions (*p*=1) yield a situation where combinations of these conditions (i.e., intermediate seasonal arrangements; 0<*p*<1) can result in competitive exclusion. This form of competitive exclusion is counterintuitive based on a linear prediction between the endpoints (*p* = 0 and 1), and represents competitive underperformance of some species. In this scenario, we assume species 2 is excluded at some point of intermediate seasonal lengths (i.e., Figure 1B). Therefore, we can focus on just the two outcomes defining stable coexistence and competitive exclusion by species 1 (Table A1). Using these criteria, we highlight what conditions allow for such a change in coexistence outcome: that is, the flip in direction of the inequality noted in red at the top of Table A1 (note that these terms define the intercepts of the approximated isoclines on the *X*_*1*_ axis; see Supplement Figure A1 for an example of the isocline geometry through this transition).

**Figure A1.**
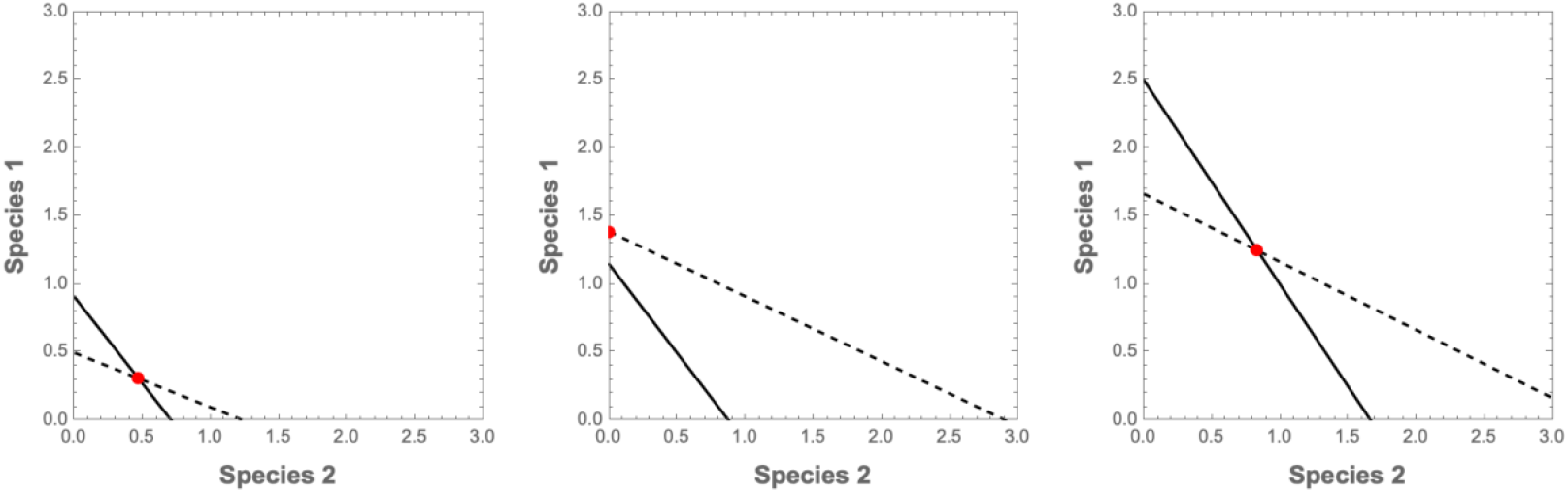
Sequence of changes in isocline geometry from *p*=0 (stable coexistence), intermediate *p* (competitive exclusion by species 1), to *p*=1 (stable coexistence). Transitions between these qualitative outcomes are seasonally-driven transcritical bifurcations.

#### Ensuring stable coexitence conditions in both seasons – defining boundary conditions

In the scenario described above, at *p*=0 we start with the following coexistence condition (the top inequality in Table A1):

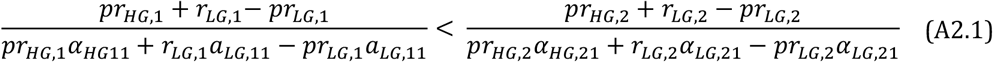

Substituting *p*=0 into (A2.1) this reduces to a form consistent with the original (non-seasonal) L-V coexistence condition:

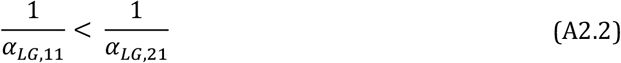

So that intraspecific competition is greater than the interspecific competition in the low-growth period as is a well-known relationship required for coexistence from classic coexistence theory.

Additionally, we note that since we are concerned with the case where both species coexist at *p*=1 (making the exclusion at intermediate *p* values counter-intuitive), we also have the following criteria:

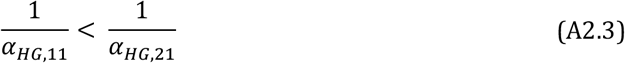

In what follows, (A2.2) and (A2.3) are our boundary conditions that define what our system approaches as *p* approaches 0 or 1 and must always be true.

#### Unpacking counterintuitive results through parameter asymmetries

We know that the above relationship (A2.1) is ultimately reversed as *p* increases such that for some given *p*, we go from the coexistence condition to the exclusion condition from Table A1, which is simply a change of sign:

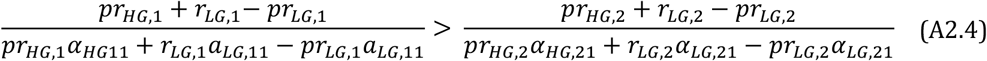

We are similarly interested in the reverse “flip” occurring, since we also have coexistence (i.e., Equation A2.1 is met) at *p*=1. Therefore, as *p* increases from 0, (A2.1) must flip to satisfy (A2.4), which must then revert back to (A2.1) as *p* approaches 1. In other words, we see two seasonally-mediated bifurcations. Since our isoclines are linear, this means that *p* must drive a nonlinear change in the attractor and we will use this when unpacking this counterintuitive (i.e., nonlinear) effect of changing *p*. The fact that *p* must cause a nonlinear path in our attractor also shows why the scenarios in Figure 1A and Figure 1B are related and the latter simply gets pushed through the boundary (i.e., *X*_*2*_=0) via a transcritical bifurcation (followed by a second transcritical bifurcation as the attractor re-enters positive state space).

Specifically, to unpack what conditions can drive this sequence of bifurcations, we require a scenario where increasing *p* (i.e., *p* grows from 0) differentially alters the LHS and RHS such that the inequality may flip (the terms in (A2.1) and (A2.4) – defining the isocline intercepts on the *X*_*1*_ axis – switch positions). Starting at *p*=0 and increasing, the sign of (A2.1) will flip when the LHS increases at a greater rate than the RHS. Similarly, when (A2.4) is satisfied and the RHS grows at a faster rate than the LHS at higher *p*, the second flip is more likely to occur.

Note that even if these terms do not fully flip, differential responses between the LHS and RHS with changing *p* will drive the same qualitative pattern in the model geometry; that is, nonlinear changes in the attractor and therefore counterintuitive changes in density (i.e., Figure 1A).

While the true response to changing *p* is governed by 8 parameters (i.e., see (A2.1) and (A2.4)), as a sketch proof we can deduce simple rules that ought to facilitate differential responses between the LHS and RHS. We can clearly see that differential responses to changing *p* will be determined by parameter asymmetries both *within* and *between* the LHS and RHS terms. Within-term differential responses will be driven by intra-specific seasonal differences in competition and growth rates, and between-term differential responses will be driven by inter-specific differences or trade-offs in competition and growth. Within these terms, responses to changing *p* will be determined by relative differences between the additive (*p*) and subtractive (-*p*) components.

To start, we can isolate the numerators to identify general parameter relationships that ought to work in the direction identified here. In this case, we can immediately see that differences in *r*_*HG,1*_ and *r*_*HG,2*_ (the positive terms multiplied by *p* in the LHS and RHS, respectively) would result in differential increases with changing *p*. Specifically, ***r***_***HG***,***1***_ **> *r***_***HG***,***2***_ ought to cause the LHS to increase at a faster rate than the RHS with increasing *p* from 0. Similarly, ***r***_***LG***,***1***_ **< *r***_***LG***,***2***_ would cause the negative part of the RHS to increase more than the LHS (thus having the same relative effect of our first condition) with increasing *p*. Both of these inter-specific growth rate differentials would act together to drive the nonlinear effect we are focusing on here and increase the potential for exclusion of species 2. We refer to these first general patterns as the **seasonal growth trade-off effect**. Note that this is especially relevant if the competition coefficients are all small (*α* < 1), since then the numerators’ responses dominate any change. Also recall that we are assuming a high-growth and low-growth period, such that *r*_*HG*_ > *r*_*LG*_ for both species, which would increase the importance of the high-growth inter-specific growth trade-offs (since they are larger and both positive terms), but this is not required.

We can use similar logic as above to deduce simple relationships that ought to drive differential responses in the denominators between the LHS and RHS of (A2.1) to changing *p* – here, largely influenced by the competition strengths (*α*′s). Due to our boundary conditions, (A2.2) and (A2.3), the denominator of the LHS will always be more influential (the *α*′s are larger) than that of the RHS in terms of the magnitude of response to changing

*p*, so *α* differences within the denominator of the LHS will largely determine if the LHS as a whole responds greater than the RHS to increasing *p*. Here, if ***α***_***LG***,**11**_ **> *α***_***HG***,**11**_ the negative part of the denominator will out-weigh the positive part, therefore causing the LHS term of (A2.1) as a whole to increase more than the RHS with increasing *p* and drive the inequality towards “flipping” to satisfy (A2.4). We can refer to this last effect as the **seasonal competition effect**.

Finally, we note that if ***α***_**11**_ ≈ ***α***_**21**_ (for both seasons), the denominators of the LHS and RHS will effectively cancel each other out and any differential responses to changing *p* will again be determined by growth rate differentials. We will refer to this as the **“sister species” effect**, since biologically similar species are likely to have similar inter- and intraspecific competition effects, and we argue this ought to amplify the result of other parameter asymmetries.

Clearly, the 8-dimensional response to changing *p* is more complex than we consider here, especially when *α*’s >> 1 (when denominators and numerators are both influential). In some instances, parameter interactions (particularly driven by the *α*’s) may mute or change the effect of any growth rate trade-offs, and in others it may be parameter combinations that contribute to the nonlinear response to *p*. For example, it is worth highlighting that if any *α* alone is large, this term will “dominate” the denominator and likely mute any asymmetries in other parameters, having a dampening effect on any differential responses between the LHS and RHS. This is likely especially true for cases where *α*_*LG*,11_ alone (being part of the constant term in the LHS denominator) is >> 1 since the denominator of the LHS would be very “heavy” and keep the LHS as a whole suppressed relative to the RHS regardless of other parameter asymmetries (i.e., changing *p* would not bring the LHS anywhere close to growing larger than the RHS).

Based on these simple mathematical observations of the relationship between parameters in the seasonal coexistence criteria (summarized in Table A1), we can identify general **rules for amplified competitive underperforming of *X***_***2***_ (i.e., nonlinear effect) over changing *p* that together would increase the potential for seasonally-mediated local extinction of species 2. These are summarized in Table A2.

**Table A2.** Conditions that are identified as amplifying competitive underperformance of species 2.

|  |  |  |
| --- | --- | --- |
| 1 | $r_{HG,1} > r_{HG,2}$ | seasonal growth trade-off effect |
| 2 | $r_{LG,1} < r_{LG,2}$ | |
| 3 | $\alpha_{LG,11} > \alpha_{HG,11}$ | seasonal competition effect |
| 4 | $\alpha_{11} \approx \alpha_{21}$ | “sister species” effect |

Note that if #3 and #4 in Table A2 hold (and keeping in mind our boundary conditions (A2.2) and (A2.3)), we can write these conditions together as, $\alpha_{LG,11} > \alpha_{LG,21} \gg \alpha_{HG,11} > \alpha_{HG,21}$ while keeping in mind that criteria #3 is more important.

Note that if #3 and #4 in Table A2 hold (and keeping in mind our boundary conditions (A2.2) and (A2.3)), we can write these conditions together as, ***α***_***LG*,11**_ > ***α***_***LG*,21**_ **>> *α***_***HG*,11**_ > ***α***_***HG***,**21**_ while keeping in mind that criteria #3 is more important.

In summarizing these relationships, note that while *α*’s can drive and/or amplify counterintuitive results themselves, they also set the boundary conditions. So, given some set of boundary conditions sufficiently far apart to allow for any nonlinear path between them (i.e., some seasonal differences in competition strength), temporal growth trade-offs between competing species have high potential for driving nonlinear effects in fluctuating environments. Specifically, temporal growth rate trade-offs have the potential to unexpectedly drive certain species to local extinction in the face of periodic environmental variation.

Note that these rules (Table A2) explain counterintuitive effects in ***one*** direction but of course this could occur in the other direction (negative nonlinear effect on species 1) if the growth trade-offs were reversed, as is the symmetric nature of this competition model. In this case, the growth trade-off effect would be amplified further by seasonal differences in *α*_22_ and *α*_12_ (the other terms defining the coexistence criteria, as shown in Table A1).

#### Numerical Results: Effects of inter versus intraspecific competition strengths on the magnitude of nonlinear effects

Figure A2 shows how changing each competition coefficient in the first two outcomes in Table A1 individually alter the magnitude of the nonlinear response (species 2 underperformance).

**Figure A2.**
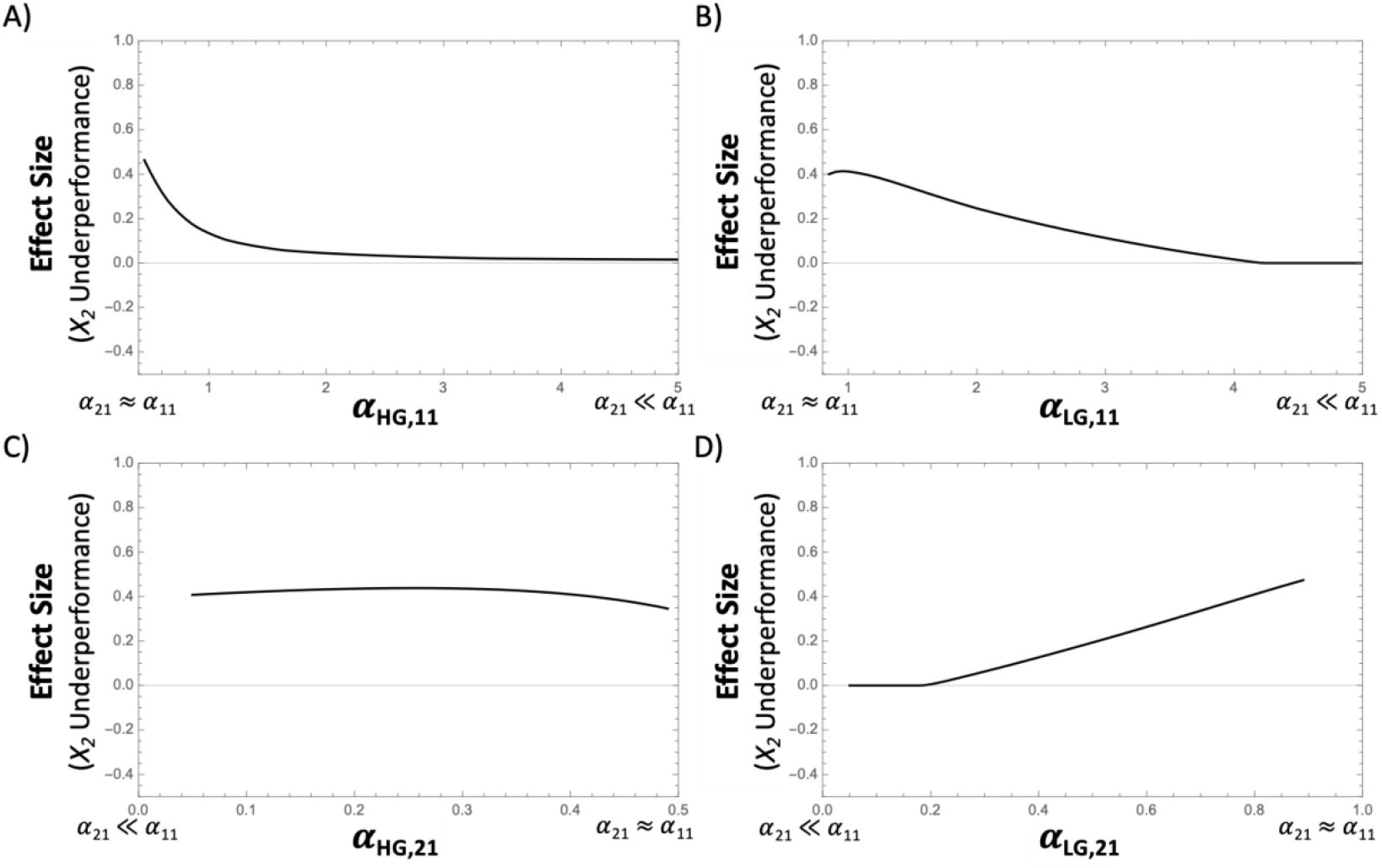
Changing *a*_11_ and *a*_21_ individually (in each season) and measuring the nonlinear effect size as the maximum difference in *X*_*2*_ over *p* = (0, 1) between the approximate equilibrium and the linear prediction based on boundary conditions.

## Notes

### Competing Interest Statement

The authors have declared no competing interest.

## References

Abbott, K. C., C. M. Heggerud, Y. C. Lai, A. Morozov, S. Petrovskii, K. Cuddington, and A. Hastings. 2024. When and why ecological systems respond to the rate rather than the magnitude of environmental changes. Biological Conservation 292:110494.

Abrams, P. A. 2022. Competition Theory in Ecology. Competition Theory in Ecology. Oxford University PressOxford.

Amarasekare, P. 2003. Competitive coexistence in spatially structured environments: A synthesis. Ecology Letters 6:1109–1122.

Arias, M. E., T. A. Cochrane, T. Piman, M. Kummu, B. S. Caruso, and T. J. Killeen. 2012. Quantifying changes in flooding and habitats in the Tonle Sap Lake (Cambodia) caused by water infrastructure development and climate change in the Mekong Basin. Journal of Environmental Management 112:53–66.

Armstrong, R. A., and R. Mcgehee. 1980. Competitive Exclusion. The American Naturalist 115:151–170.

Bartley, T. J., K. S. McCann, C. Bieg, K. Cazelles, M. Granados, M. M. Guzzo, A. S. MacDougall, et al. 2019. Food web rewiring in a changing world. Nature Ecology & Evolution 3:345–354.

Bieg, C., G. Gellner, and K. S. McCann. 2023. Stability of consumer-resource interactions in periodic environments. Proceedings of the Royal Society B: Biological Sciences 290.

Bieg, C., H. Vallès, A. Tewfik, B. E. Lapointe, and K. S. McCann. 2022. Towards a multistressor theory for coral reefs in a changing world. bioRxiv.

Brown, J. S. 1989. Desert Rodent Community Structure: A Test of Four Mechanisms of Coexistence. Ecological Monographs 59:1–20.

Cáceres, C. E. 1997. Temporal variation, dormancy, and coexistence: A field test of the storage effect. Proceedings of the National Academy of Sciences of the United States of America 94:9171–9175.

Chesson, P. 2018. Updates on mechanisms of maintenance of species diversity. Journal of Ecology 106:1773–1794.

Chesson, P., R. L. E. Gebauer, S. Schwinning, N. Huntly, K. Wiegand, M. S. K. Ernest, A. Sher, et al. 2004. Resource pulses, species interactions, and diversity maintenance in arid and semiarid environments. Oecologia 141:236–253.

Chesson, P., and N. Huntly. 1997. The Roles of Harsh and Fluctuating Conditions in the Dynamics of Ecological Communities. The American Naturalist 150:730–757.

Cushing, J. M. 1980. Two species competition in a periodic environment. Journal of Mathematical Biology 10:385–400.

Dillon, M. E., H. A. Woods, G. Wang, S. B. Fey, D. A. Vasseur, R. S. Telemeco, K. Marshall, et al. 2016. Life in the Frequency Domain: the Biological Impacts of Changes in Climate Variability at Multiple Time Scales. Integrative and Comparative Biology 56:14–30.

Greyson-Gaito, C. J., G. Gellner, and K. S. McCann. 2023. Life-history speed, population disappearances and noise-induced ratchet effects. Proceedings of the Royal Society B: Biological Sciences 290.

Hadeler, K. P., and T. Hillen. 2007. Coupled dynamics and quiescent phases. Math Everywhere: Deterministic and Stochastic Modelling in Biomedicine, Economics and Industry. Dedicated to the 60th Birthday of Vincenzo Capasso 7–23.

Harabiš, F., A. Dolný, and J. Šipoš. 2012. Enigmatic adult overwintering in damselflies: coexistence as weaker intraguild competitors due to niche separation in time. Population Ecology 54:549–556.

Hastings, A., K. C. Abbott, K. Cuddington, T. B. Francis, Y. C. Lai, A. Morozov, S. Petrovskii, et al. 2021. Effects of stochasticity on the length and behaviour of ecological transients. Journal of the Royal Society Interface 18.

Hastings, A., K. C. Abbott, K. Cuddington, T. Francis, G. Gellner, Y. C. Lai, A. Morozov, et al. 2018. Transient phenomena in ecology. Science 361.

Lancaster, L. T., G. Morrison, and R. N. Fitt. 2017. Life history trade-offs, the intensity of competition, and coexistence in novel and evolving communities under climate change. Philosophical Transactions of the Royal Society B: Biological Sciences 372:20160046.

Li, L., and P. Chesson. 2016. The effects of dynamical rates on species coexistence in a variable environment: The paradox of the plankton revisited. American Naturalist 188:E46–E58.

Litchman, E., and C. A. Klausmeier. 2001. Competition of phytoplankton under fluctuating light. American Naturalist 157:170–187.

Mathias, A., and P. Chesson. 2013. Coexistence and evolutionary dynamics mediated by seasonal environmental variation in annual plant communities. Theoretical Population Biology 84:56–71.

McMeans, B. C., K. S. McCann, M. M. Guzzo, T. J. Bartley, C. Bieg, P. J. Blanchfield, T. Fernandes, et al. 2020. Winter in water: differential responses and the maintenance of biodiversity. Ecology Letters 23:922–938.

McMeans, B. C., K. S. McCann, T. D. Tunney, A. T. Fisk, A. M. Muir, N. Lester, B. Shuter, et al. 2016. The adaptive capacity of lake food webs: from individuals to ecosystems. Ecological Monographs 86:4–19.

Morozov, A., U. Feudel, A. Hastings, K. C. Abbott, K. Cuddington, C. M. Heggerud, and S. Petrovskii. 2024. Long-living transients in ecological models: Recent progress, new challenges, and open questions. Physics of Life Reviews 51:423–441.

Mougi, A. 2020. Polyrhythmic foraging and competitive coexistence. Scientific Reports 10:20282.

Namba, T. 1984. Competitive Co-existence in a seasonally fluctuating environment. Journal of Theoretical Biology 111:369–386.

Namba, T., and S. Takahashi. 1993. Competitive Coexistence in a Seasonally Fluctuating Environment II. Multiple Stable States and Invasion Success. Theoretical Population Biology 44:374–402.

Neves, M. P., P. Kratina, R. L. Delariva, J. I. Jones, and C. B. Fialho. 2021. Seasonal feeding plasticity can facilitate coexistence of dominant omnivores in Neotropical streams. Reviews in Fish Biology and Fisheries 31:417–432.

Penk, M. R., J. M. Jeschke, D. Minchin, and I. Donohue. 2016. Warming can enhance invasion success through asymmetries in energetic performance. (B. Griffen, ed.) Journal of Animal Ecology 85:419–426.

Schoener, T. W. 1982. The controversy over interspecific competition. American Scientist 70:586–595.

Schreiber, S. J. 2021. Positively and negatively autocorrelated environmental fluctuations have opposing effects on species coexistence. American Naturalist 197:405–414.

Scott, A. M., C. Bieg, B. C. McMeans, and K. S. McCann. 2023. Coexistence in Periodic Environments. bioRxiv.

Scranton, K., and D. A. Vasseur. 2016. Coexistence and emergent neutrality generate synchrony among competitors in fluctuating environments. Theoretical Ecology 9:353–363.

Sharma, S., K. Blagrave, J. J. Magnuson, C. M. O’Reilly, S. Oliver, R. D. Batt, M. R. Magee, et al. 2019. Widespread loss of lake ice around the Northern Hemisphere in a warming world. Nature Climate Change 9:227–231.

Vasseur, D. A., and P. Yodzis. 2004. The color of environmental noise. Ecology 85:1146–1152.

White, E. R., and A. Hastings. 2020. Seasonality in ecology: Progress and prospects in heory. Ecological Complexity 44:100867.

Woolway, R. I., S. Sharma, G. A. Weyhenmeyer, A. Debolskiy, M. Golub, D. Mercado-Bettín, M. Perroud, et al. 2021. Phenological shifts in lake stratification under climate change. Nature Communications 12:2318.

Yeh, S. W., J. S. Kug, B. Dewitte, M. H. Kwon, B. P. Kirtman, and F. F. Jin. 2009. El Nĩo in a changing climate. Nature 461:511–514.

## References

Scott, A. M., C. Bieg, B. C. McMeans, and K. S. McCann. 2023. Coexistence in Periodic Environments. bioRxiv 2023.02.24.529749.

